# Degradation of key photosynthetic genes in the critically endangered semi-aquatic flowering plant *Saniculiphyllum guangxiense* (Saxifragaceae)

**DOI:** 10.1101/2019.12.22.886283

**Authors:** Ryan A. Folk, Neeka Sewnath, Chun-Lei Xiang, Brandon T. Sinn, Robert P. Guralnick

## Abstract

**Background:** Plastid gene loss and pseudogenization has been widely documented in parasitic and mycoheterotrophic plants, which have relaxed selective constraints on photosynthetic function. More enigmatic are sporadic reports of degradation and loss of important photosynthesis genes in lineages thought to be fully photosynthetic. Here we report the complete plastid genome of *Saniculiphyllum guangxiense*, a critically endangered and phylogenetically isolated plant lineage, along with genomic evidence of reduced chloroplast function. We also report 22 additional plastid genomes representing the diversity of its containing clade Saxifragales, characterizing gene content and placing variation in a broader phylogenetic context.

**Results:** We find that the plastid genome of *Saniculiphyllum* has experienced pseudogenization of five genes of the NDH complex (*ndhA*, *ndhB, ndhD*, *ndhF*, and *ndhK*), previously reported in flowering plants with an aquatic habit, as well as the more surprising pseudogenization of two genes more central to photosynthesis (*ccsA* and *cemA*), contrasting with strong phylogenetic conservatism of plastid gene content in all other sampled Saxifragales. These genes participate in photooxidative protection, cytochrome synthesis, and carbon uptake. Nuclear paralogs exist for all seven plastid pseudogenes, yet these are also unlikely to be functional.

**Conclusions:** *Saniculiphyllum* appears to represent the greatest degree of plastid gene loss observed to date in any fully photosynthetic lineage, yet plastid genome length, structure, and substitution rate are within the variation previously reported for photosynthetic plants. These results highlight the increasingly appreciated dynamism of plastid genomes, otherwise highly conserved across a billion years of green plant evolution, in plants with highly specialized life history traits.

## Background

Plastid genome structure and content is highly conserved among most of the ∼500,000 species of land plants and their closest green algal relatives. Nevertheless, widespread loss or pseudogenization of photosynthetic genes is a familiar feature of the plastids of diverse non-photosynthetic plant lineages, reflecting the reduced need for photosynthetic genes in lineages with heterotrophic strategies. Accumulating evidence, however, has increasingly documented the loss of “accessory” photosynthetic genes, only conditionally essential under stress, in fully photosynthetic plants. Although not universal, many of these losses are associated with highly specialized life history traits such as aquatic habit [1–3], carnivory [4, 5], and a mycoheterotrophic life-stage [6]; the functional significance of these losses remains enigmatic [7].

*Saniculiphyllum guangxiense* C.Y. Wu & T.C. Ku is a semi-aquatic flowering plant now restricted to a miniscule area in Yunnan province, China. It grows partially submersed in the flow of small shaded waterfalls, and is critically endangered, with only four small extant populations in an area ∼10 km^2^ known to science, as well as several other populations known to have been extirpated within the last 30 years [8]. Consistent with the isolated morphological and ecological traits of this lineage within the family Saxifragaceae, its phylogenetic affinities remain uncertain. The most recent attempts to place this species [8–10] exhibit strong disagreement. [8], using six loci generated by Sanger sequencing, could not confidently place this lineage beyond its membership in the Heucheroid clade, while [9], using the same genetic loci, were able to place this lineage with 0.93-1.0 posterior probability (depending on the analysis) as sister to the *Boykinia* group, a difference Deng et al. attribute to alignment differences in a single rapidly evolving genetic locus (ITS). Relationships in these studies based on Sanger sequencing data differ substantially in several areas from those recovered on the basis of more than 300 nuclear genes [10], where *Saniculiphyllum* was placed with moderate bootstrap support (80%) as sister to a clade containing the *Astilbe* and *Boykinia* groups.

In the course of organellar genome surveys across Saxifragales, we found anomalous photosynthetic gene sequences in *Saniculiphyllum.* Here, we report new plastid genome sequences of phylogenetically pivotal taxa, analyze plastid gene evolution across the Saxifragales and place the *Saniculiphyllum* plastid genome in a phylogenetic context to assess evolutionary relationships and rates of plastid evolution.

## Results

### Assembly results

For all samples, NOVOPlasty successfully assembled a complete circular genome. We individually confirmed all sequence features noted below by mapping the reads back to the assembly, and found no evidence of misassembly.

### Basic genome features

*Saniculiphyllum* has a chloroplast genome 151,704 bp long (Fig. 1). The large-scale structure of the genome is canonical for land plants, with an inverted repeat (26,109 bp) separating the large-single-copy region (LSC; 84,479 bp) and small-single-copy region (SSC, 15,007 bp). Excluding putative pseudogenes, gene content was as expected, comprising 73 distinct protein-coding genes, 30 tRNA genes, and 4 rRNA genes.

**Figure 1.**
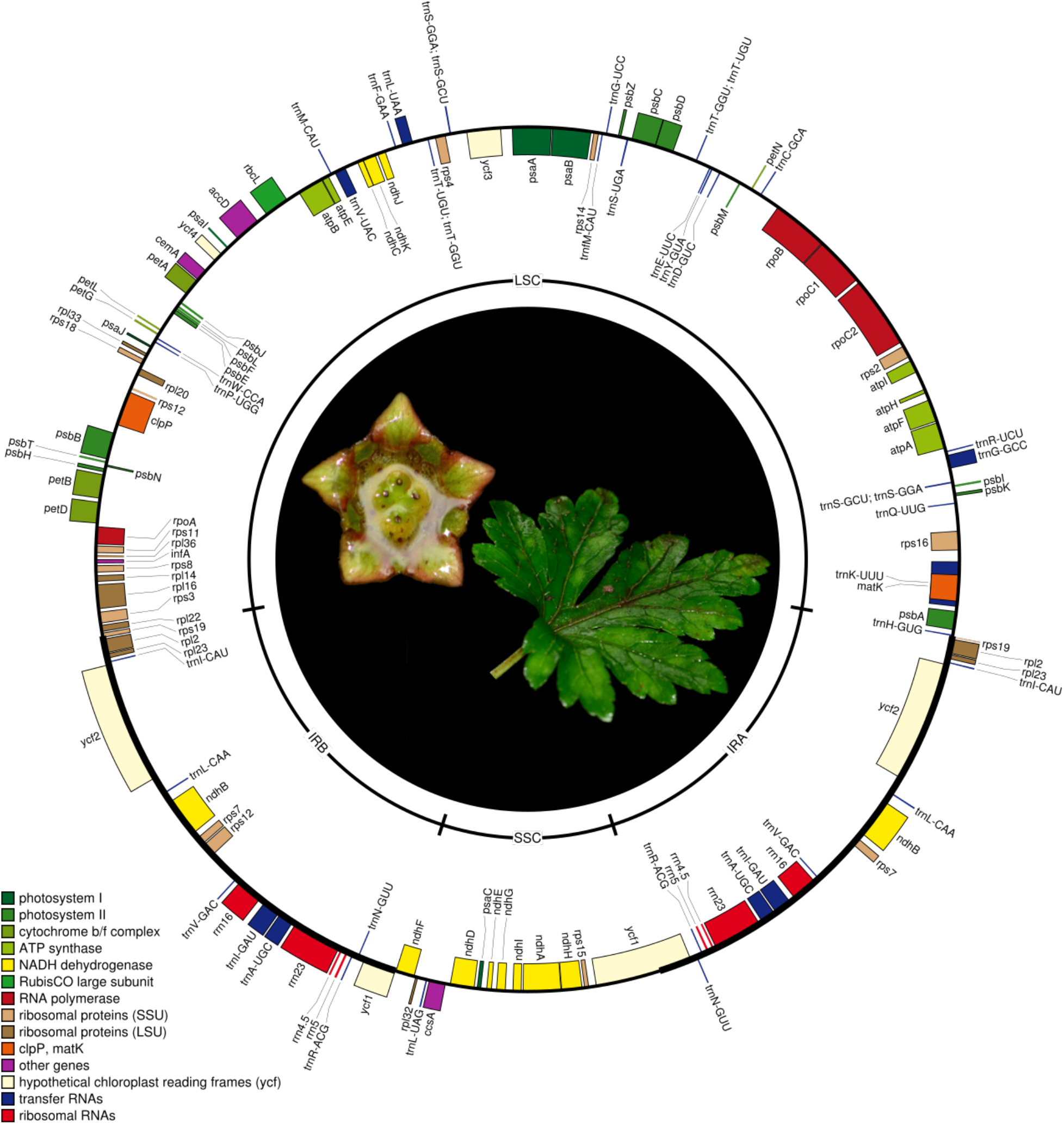
Gene map of the *Saniculiphyllum* plastome built using OrganellarGenomeDRAW [11]; genes marked on the outside face of the circle are transcribed counter-clockwise and those inside the circle are transcribed clockwise. Center photo: *Saniculiphyllum* flower and leaf; photo credit: C.-L. X.

### Evidence for pseudogenization

We found genomic evidence for pseudogenization in 5 genes of the NDH complex (*ndhA, ndhB, ndhD, ndhF*, and *ndhK*), and two other photosynthetic genes (*cemA, ccsA*), summarized in Table 1. These were either driven by frame-shift mutations (*ccsA, ndhA, ndhD*, and *ndhF*) or by premature stop codons without a frameshift (due to a point mutation in *ndhB* and a short inversion in *ndhK*). Three genes (*cemA*, *ndhD*, and *ndhF*) lack much of the conserved gene sequence due to large deletions >100 bp. Among these, *cemA* has no premature stop codons, but it has an unconventional predicted protein size (5 extra amino acids) in a gene that otherwise shows no size variation in Saxifragales; while lacking 18% of the 3’ end of this gene, *Saniculiphyllum* has 137 additional bp before a novel stop codon, the sequence of which is homologous with adjacent intergenic spacers in its relatives, making it unlikely that this sequence is functional. Additionally, frameshift has resulted in the loss of the conserved stop codon site of *ndhA*. The three genes with large deletions (*cemA*, *ndhD*, and *ndhF*) also have hydrophobicity outside the range of variation of other Saxifragales (*cemA* 50% hydrophobic amino acids vs. the 95% confidence interval for other Saxifragales [50.4%, 52.2%]; *ndhD* 47.8% vs. [62.2%, 63.6%]; *ndhF* 54.9% vs. [55.6%, 58.2%]).

**Table 1.**
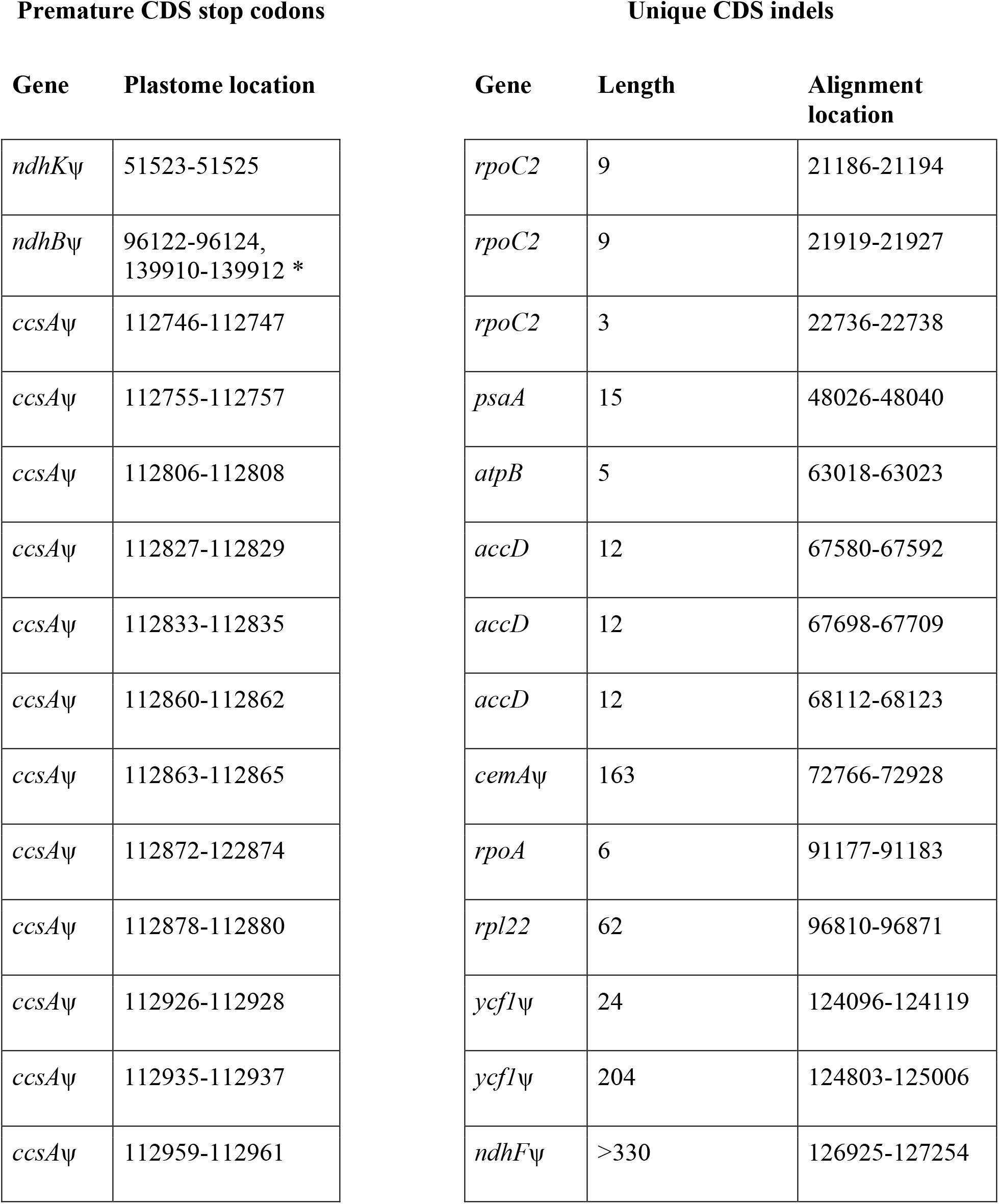
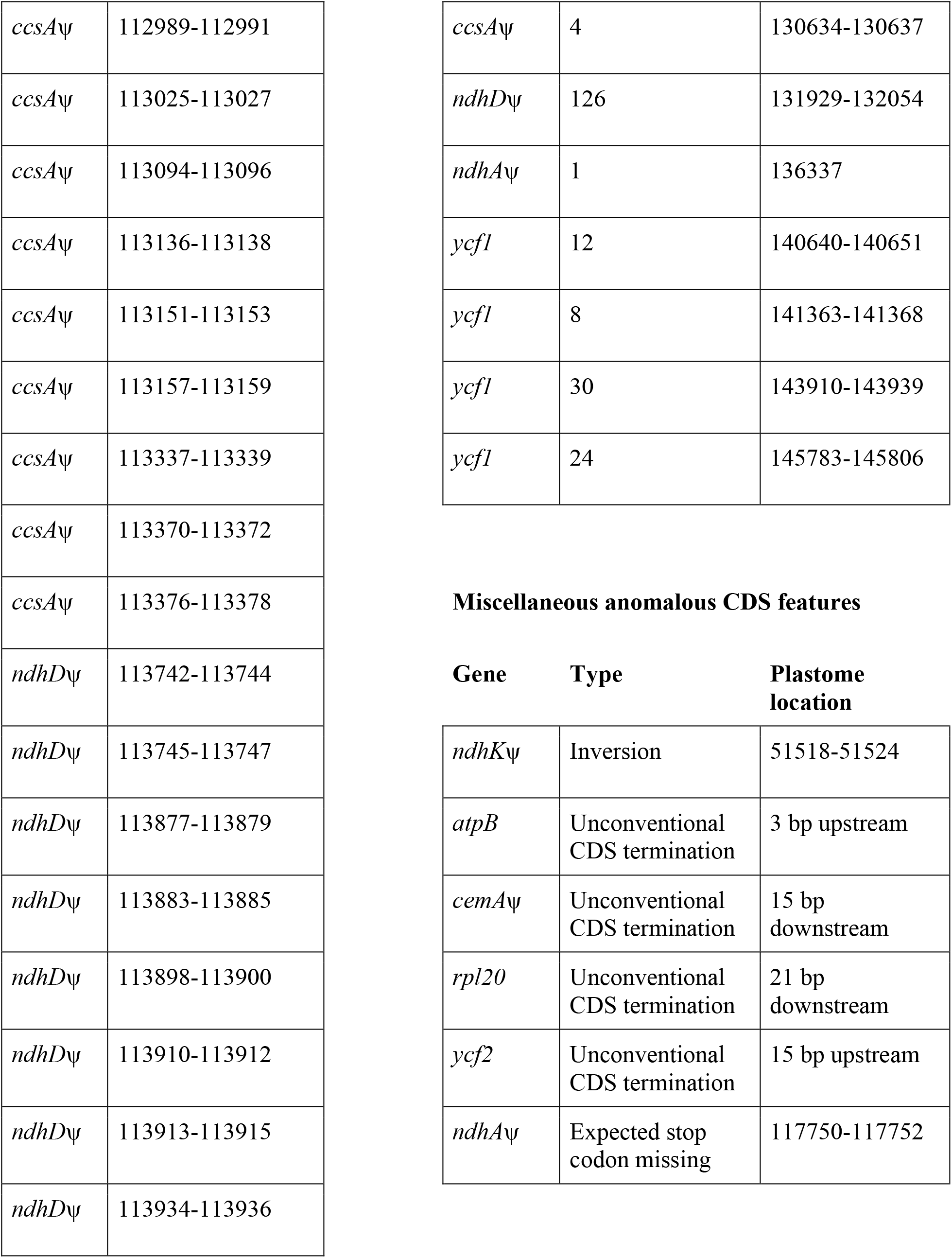
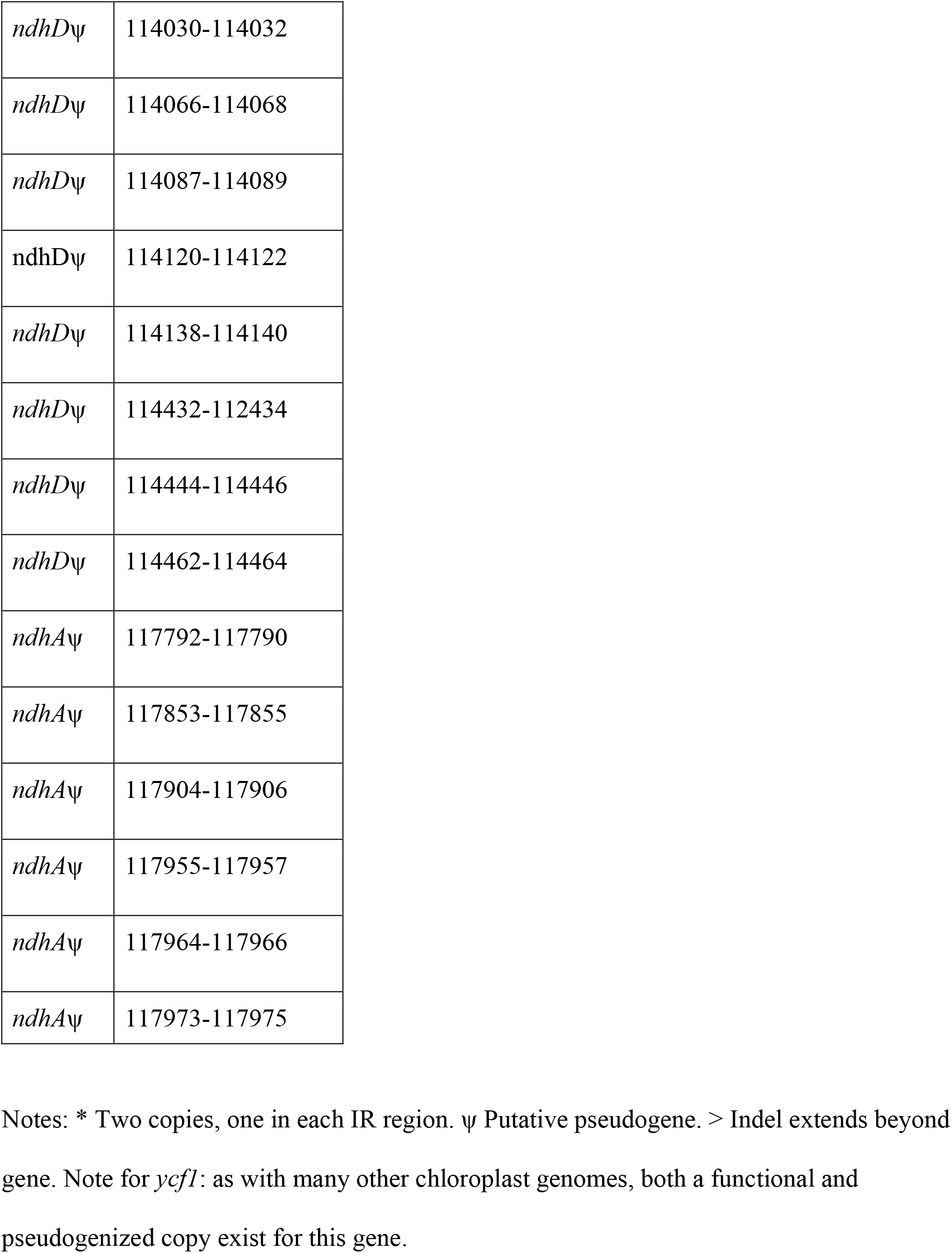
Summary of premature stop codons, large/frame-shifting indels, and other anomalous genome features unique to *Saniculiphyllum*.

| Premature CDS stop codons |  | Unique CDS indels |  |  |
| --- | --- | --- | --- | --- |
| Gene | Plastome location | Gene | Length | Alignment location |
| *ndhK*ψ | 51523-51525 | *rpoC2* | 9 | 21186-21194 |
| *ndhB*ψ | 96122-96124,<br>139910-139912 * | *rpoC2* | 9 | 21919-21927 |
| *ccsA*ψ | 112746-112747 | *rpoC2* | 3 | 22736-22738 |
| *ccsA*ψ | 112755-112757 | *psaA* | 15 | 48026-48040 |
| *ccsA*ψ | 112806-112808 | *atpB* | 5 | 63018-63023 |
| *ccsA*ψ | 112827-112829 | *accD* | 12 | 67580-67592 |
| *ccsA*ψ | 112833-112835 | *accD* | 12 | 67698-67709 |
| *ccsA*ψ | 112860-112862 | *accD* | 12 | 68112-68123 |
| *ccsA*ψ | 112863-112865 | *cemA*ψ | 163 | 72766-72928 |
| *ccsA*ψ | 112872-122874 | *rpoA* | 6 | 91177-91183 |
| *ccsA*ψ | 112878-112880 | *rpl22* | 62 | 96810-96871 |
| *ccsA*ψ | 112926-112928 | *ycf1*ψ | 24 | 124096-124119 |
| *ccsA*ψ | 112935-112937 | *ycf1*ψ | 204 | 124803-125006 |
| *ccsA*ψ | 112959-112961 | *ndhF*ψ | >330 | 126925-127254 |

|  |  |
| --- | --- |
| *ccsA*ψ | 112989-112991 |
| *ccsA*ψ | 113025-113027 |
| *ccsA*ψ | 113094-113096 |
| *ccsA*ψ | 113136-113138 |
| *ccsA*ψ | 113151-113153 |
| *ccsA*ψ | 113157-113159 |
| *ccsA*ψ | 113337-113339 |
| *ccsA*ψ | 113370-113372 |
| *ccsA*ψ | 113376-113378 |
| *ndhD*ψ | 113742-113744 |
| *ndhD*ψ | 113745-113747 |
| *ndhD*ψ | 113877-113879 |
| *ndhD*ψ | 113883-113885 |
| *ndhD*ψ | 113898-113900 |
| *ndhD*ψ | 113910-113912 |
| *ndhD*ψ | 113913-113915 |
| *ndhD*ψ | 113934-113936 |

|  |  |  |
| --- | --- | --- |
| *ccsA*ψ | 4 | 130634-130637 |
| *ndhD*ψ | 126 | 131929-132054 |
| *ndhA*ψ | 1 | 136337 |
| *ycf1* | 12 | 140640-140651 |
| *ycf1* | 8 | 141363-141368 |
| *ycf1* | 30 | 143910-143939 |
| *ycf1* | 24 | 145783-145806 |

**Gene Type Plastome location**
|  |  |  |
| --- | --- | --- |
| *ndhK*ψ | Inversion | 51518-51524 |
| *atpB* | Unconventional CDS termination | 3 bp upstream |
| *cemA*ψ | Unconventional CDS termination | 15 bp downstream |
| *rpl20* | Unconventional CDS termination | 21 bp downstream |
| *ycf2* | Unconventional CDS termination | 15 bp upstream |
| *ndhA*ψ | Expected stop codon missing | 117750-117752 |

|  |  |
| --- | --- |
| *ndhD*ψ | 114030-114032 |
| *ndhD*ψ | 114066-114068 |
| *ndhD*ψ | 114087-114089 |
| *ndhD*ψ | 114120-114122 |
| *ndhD*ψ | 114138-114140 |
| *ndhD*ψ | 114432-114434 |
| *ndhD*ψ | 114444-114446 |
| *ndhD*ψ | 114462-114464 |
| *ndhA*ψ | 117792-117790 |
| *ndhA*ψ | 117853-117855 |
| *ndhA*ψ | 117904-117906 |
| *ndhA*ψ | 117955-117957 |
| *ndhA*ψ | 117964-117966 |
| *ndhA*ψ | 117973-117975 |

Notes: \* Two copies, one in each IR region. ψ Putative pseudogene. > Indel extends beyond gene. Note for *ycfI*: as with many other chloroplast genomes, both a functional and pseudogenized copy exist for this gene.

### Evidence for paralogs of pseudogenes

For the three genes with large deletions (*cemA*, *ndhD*, and *ndhF*), we used the *Leptarrhena* sequence for the missing DNA to probe for potential nuclear or mitochondrial paralogs that could be functional; otherwise we used the entire CDS of this taxon. For all seven novel pseudogenes, we found evidence of paralogs outside of the assembled chloroplast genome, some of which are more conserved in sequence and lack the anomalous features of plastid pseudogenes (Supplementary Figs. S1-7). This includes copies of *cemA*, *ndhD*, and *ndhF* without the large deletions found in the plastid copy. However, with the exception of partial assembled sequences of *ndhF*, these paralogs all have either the same premature stop codons of the plastid copy or novel premature stop codons, and are also unlikely to be functional. These paralogs likely originate in the nucleus on the basis of sequence coverage, which was orders of magnitude lower (SPAdes calculated kmer coverage ∼1-5X) than that expected for either the plastid or the mitochondrion (kmer coverage 100-2000X).

With the exception of *ndhK*, where we recovered 4 independent lineages of *Saniculiphyllum* paralogs, gene genealogies (Figs. S1-7) were consistent with a recent origin of paralogs of the seven pseudogenes. In the *ccsA* gene genealogy, the *Saxifraga stolonifera* Curtis plastid ortholog was placed within a *Saniculiphyllum* clade without support, but otherwise (*cemA, ndhA, ndhB, ndhD, ndhF*) the *Saniculiphyllum* paralogs were recovered as monophyletic.

### Other anomalous features

Several genes show slight variations in within-frame start and stop codon positions in Saxifragales, but *Saniculiphyllum* shows more variation than any other species we sampled, with four genes showing unique CDS terminations (*atpB, cemA, rpl20, ycf2*; Table 1), of which none but *rpl20* show any size variation in other Saxifragales species. While still within the typical length of photosynthetic plastid genomes, *Saniculiphyllum* was significantly smaller than the mean for Saxifragales species (one-tailed t-test, *p* = 1.485e-10).

Interestingly, the percent of total genomic DNA from the plastid genome was also significantly smaller in *Saniculiphyllum* (3.4%) compared to other Saxifragales (one-tailed t-test, *p* = 1.629e-07); the mean of our Saxifragales species sampled here was 10.1%, identical to a mean of 10.1% recovered with further Saxifragaceae species sampled in [12]).

### Phylogenetic analysis

The plastome alignment length was 172,773 bp, with 9.9% of the alignment comprising gap characters, and 38,332 parsimony-informative characters excluding the gap characters. Backbone relationships in the chloroplast genome phylogeny were congruent with [10] (Fig. 2). Although receiving maximal bootstrap support, the placement of *Saniculiphyllum* we recovered is different from all previous efforts to place this taxon, none of which agree among themselves and none of which achieved greater than moderate support [8–10]. Our placement resembles [9, 10] in placing *Saniculiphyllum* in a clade comprising the *Astilbe* Buch.-Ham., *Boykinia* Raf., and *Leptarrhena* groups, but the novel placement reported here is sister to *Leptarrhena.* Despite its divergent plastome features, genome-wide substitution rates are not elevated in *Saniculiphyllum* (Fig. 2).

**Figure 2.**
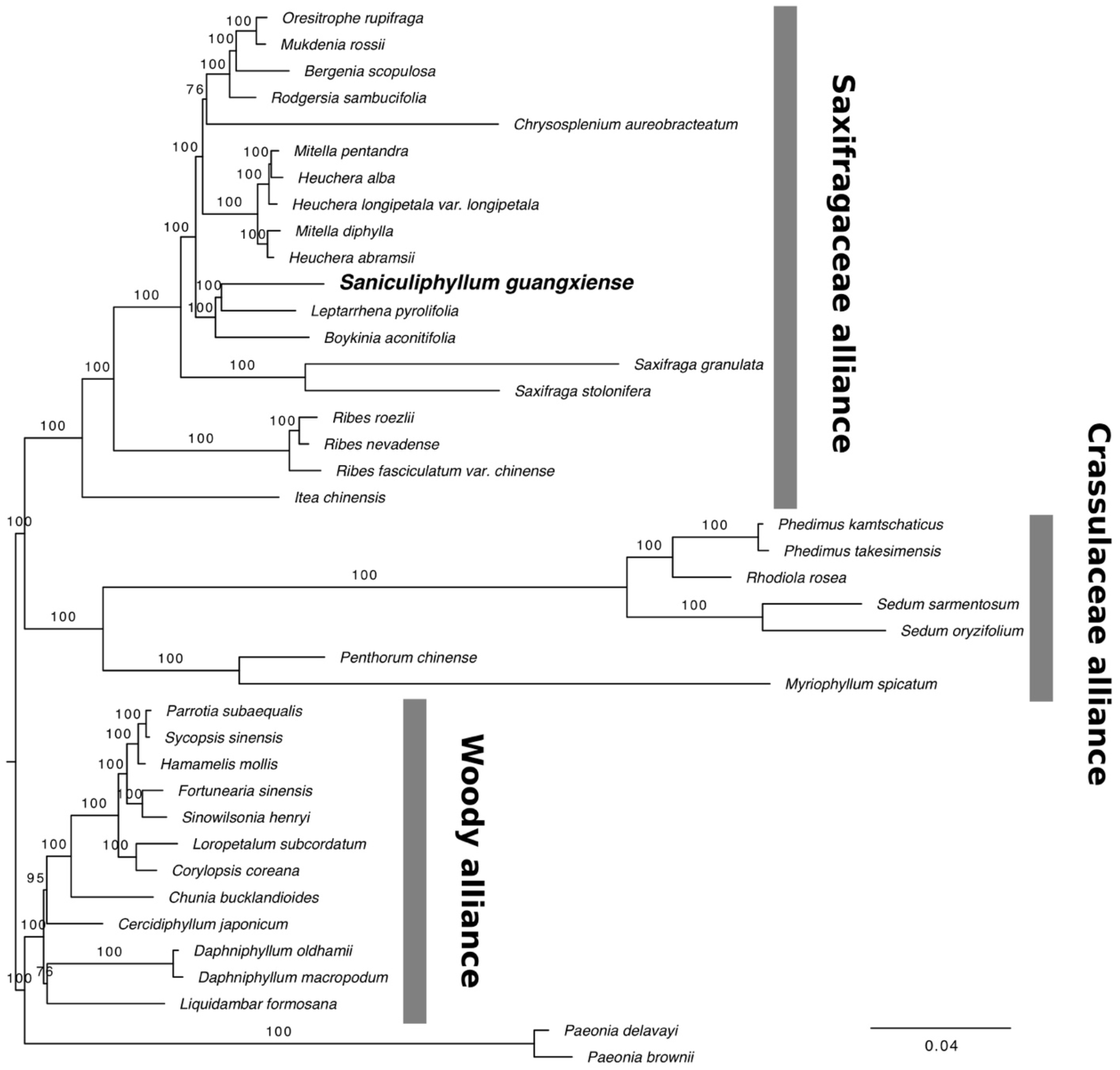
ML phylogeny of Saxifragales plastid genomes. *Saniculiphyllum* shown in bold; labelled clades correspond to the terminology of [13]. Branch labels represent bootstrap frequencies.

## Discussion

### Gene loss

In total, we found genomic evidence for seven putative pseudogenes in the *Saniculiphyllum* plastid genome. Five of these (*ndhA, ndhB, ndhD, ndhF*, and *ndhK*), are genes of the NDH complex. These genes are highly conserved across the land plants and related green algae [7]. Most losses of plastid gene function have been associated with parasitic and mycoheterotrophic plants, which presumably have few functional constraints on photosynthetic gene evolution. Degradation of genes in the NDH complex has nevertheless been observed in several fully photosynthetic lineages with a variety of life history traits: woody perennials in Pinaceae and Gnetales (both gymnosperms), short-lived perennials in Geraniaceae (eudicots: rosids), carnivorous and often aquatic plants of Lentibulariaceae (eudicots: asterids), various photosynthetic members of Orchidaceae (monocot), and aquatic members of Alismatales (monocot) and Podostemataceae (rosid; [1, 3, 6, 7, 14–17]). The primary function of the NDH complex is thought to be reduction of photooxidative stress under fluctuating light conditions. While the NDH complex appears dispensable under mild growth conditions [18], experimental evidence from knockouts of single *ndh* genes shows that a complete and intact complex is essential for efficient photosynthesis and robust plant growth under stressful conditions [14].

More unusual than loss of NDH function is the clear pseudogenization of two other photosynthesis-specific genes, for which we report the first absence in a fully photosynthetic plant. The gene *cemA* encodes a protein involved in carbon uptake; while not essential for photosynthesis, photosynthetic efficiency is reduced under high light environments in *Chlamydomonas* Ehrenb. mutants lacking this gene [19]. The gene *ccsA* encodes a protein involved in heme attachment to chloroplast cytochrome c [20]. *ccsA*, at least in *Chlamydomonas*, is essential for System II photosynthesis [20]. Both *cemA* and *ccsA* are conserved across primary photosynthetic eukaryotes and even cyanobacteria [19, 21].

### Evidence for paralogs in the nucleus

We successfully found and assembled paralogs for all seven novel putative chloroplast pseudogenes in *Saniculiphyllum.* Many of these paralogs are of more conserved sequence than that of the assembled plastid genome; with the exception of *ndhK* these appear to have originated primarily after the divergence of *Saniculiphyllum* from other Saxifragaceae lineages. On the basis of coverage, these are likely to represent NUPTs (nuclear sequences of plastid origin; [22]). While we do not have direct evidence for functional importation of a functional photosynthetic protein from these paralogs into the chloroplast, and indeed most of them show signs of pseudogenization, our results are consistent with growing evidence of a slow transfer of organellar gene content into nuclear genomes [22, 23], a process associated with frequent non-homologous recombinational repair between these genomes [24].

### Other genome anomalies

We also observed unusual CDS terminations upstream or downstream of closely related Saxifragales plastid genomes in four genes; these do not result in frameshifts but expected protein product are of unexpected length. Although less dramatic than the pseudogenization patterns we observed, the lack of length conservation in *Saniculiphyllum* is markedly greater compared to close relatives. Likewise, while the *Saniculiphyllum* plastome is far longer than many non-photosynthetic plants (reviewed in [25]), it is among the shortest in Saxifragales due to large deletions in coding and non-coding regions throughout the plastome.

Despite having one of the most divergent plastid genomes in Saxifragales, there is no evidence for elevated substitution rates in *Saniculiphyllum* based on phylogenetic branch length estimated from the entire plastid genome (Fig. 2). Likewise, we implemented tests on dN/dS ratios in the seven putative pseudogenes, demonstrating that *Saniculiphyllum* does not show significantly different selection regimes at the codon level compared to related lineages (all *p* > 0.05; dN/dS < 1 in all cases with mean 0.0319). These results suggest that *Saniculiphyllum* primarily differs in its plastid genome evolution via deletions and rare novel stop codons without any detectable global relaxation of purifying selection at the codon level. Dosage of plastid DNA relative to the nucleus also appears to be low in *Saniculiphyllum* compared to relatives, likely representing either a reduction in plastids per cell or a reduction in genome copy number per plastid.

### Evolutionary relationships

This work also represents the first robust phylogenomic placement of *Saniculiphyllum*, an important group for interpreting morphological evolution in Saxifragaceae [8]. We confirm a close relationship with the *Boykinia* and *Leptarrhena* groups, with which it shares axile placentation, determinate cymose inflorescences, and a strongly rhizomatous habit. However, representatives of the *Astilbe* group and several others have yet to be sampled; denser taxon sampling is needed to confirm the placement reported here.

## Conclusions

Although chloroplast genome evolution in Saxifragales has been previously understood as very conservative [26], further sampling has revealed surprising plastid variation in one of its rarest and most unusual lineages. Similar but less extreme patterns of gene loss have been observed before in aquatic members of order Alismatales and Podostemaceae, and appear to represent multiple independent evolutionary events [1, 3], suggesting a possible relationship with life history. Nevertheless, this putative correlation is imperfect; unlike the partly aerial *Saniculiphyllum*, Alismatales contains some of the most thoroughly aquatic-adapted angiosperms, including the only examples of aquatic pollination [1]. By contrast, *Myriophyllum*, a completely aquatic Saxifragales lineage, shows conventional gene content [27], as do many other aquatic plastid genomes (e.g., *Nelumbo* Adans. [28], *Nymphaea* L. [29], *Lemna* L. [30]).

It is tempting to speculate on the relationship between loss of photosynthetic gene content and the imperiled conservation status of *Saniculiphyllum*. Unfortunately, we understand little of the functional significance of plastid gene content outside of model organisms, highlighting the need for characterization of plastid genomes and further examination of the relationship between organellar genome evolution and life history traits.

## Methods

### Sampling

We sequenced 23 plastomes in total to increase phylogenetic representation. Other than *Saniculiphyllum*, we sampled 16 further taxa of Saxifragaceae to cover most of the major recognized clades recognized in [9], and six further Saxifragales outgroups to increase representation in the woody alliance (cf. [13]).

### DNA extraction and sequencing

Whole genomic DNAs were isolated from fresh or silica-dried leaf material using a modified CTAB extraction protocol [31]. Taxa were chosen to represent lineages across Saxifragales. Sequencing was performed either at RAPiD Genomics (Gainesville, Florida, U.S.A.) with 150 bp paired-end Illumina HiSeq sequencing or with 100 bp paired-end BGISEQ-500 sequencing at BGI (Shenzhen, Guangdong, P.R. China), in both cases with an insert size of approximately 300 bp (summarized in Table 2).

**Table 2.**
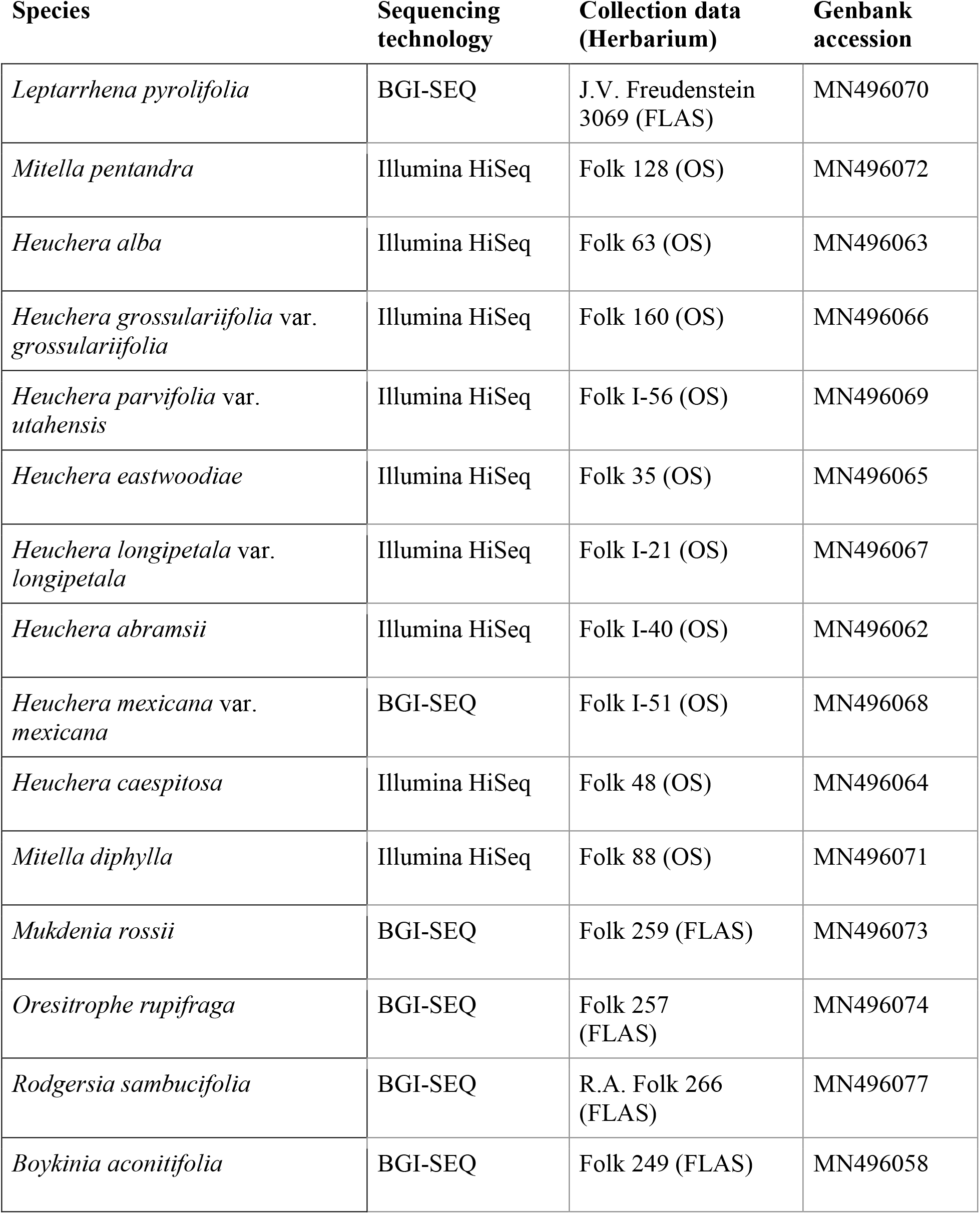
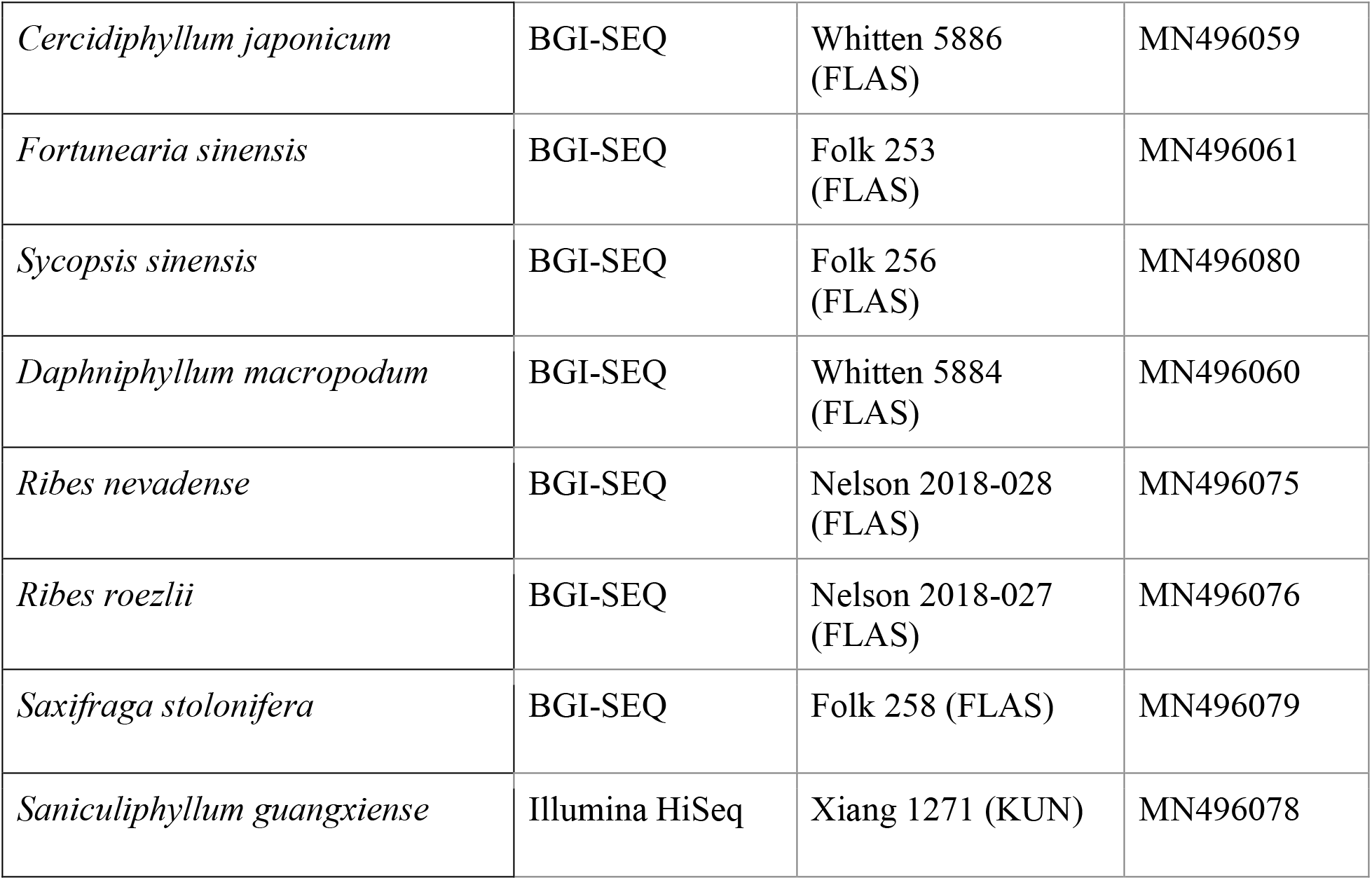
Summary of new chloroplast genome sequences reported in this paper.

### Genome assembly

We used NOVOPlasty v. 3.2 [32] to assemble chloroplast genomes for all sequenced taxa. For each sample, we ran two assemblies using *rbcL* and *matK* seed reference genes from the plastid genome of *Heuchera parviflora* var. *saurensis* R.A. Folk [12]. Reads were not quality filtered following developer recommendations. We have found that NOVOPlasty assemblies can be negatively affected by very large short read datasets; datasets were normalized to 8 million raw reads per sample for HiSeq data and 4 million for BGI-SEQ samples (∼100-500X plastid coverage). The orientation of the small-single copy region relative to the rest of the genome was manually standardized across samples.

Annotations were performed in Geneious R9 using the *Heuchera* reference plastid genome and a cutoff of 70% sequence identity, and draft annotated plastid genomes were aligned and manually examined for annotation accuracy. Additionally, all premature stop codons, inversions, frameshifting indels, and other unusual features were individually verified visually by mapping the original reads back to the assembled plastid genomes using the Geneious read mapping algorithm [33]. We also calculated the percent of chloroplast sequences in the total DNA from these mapped reads using SAMtools [34].

For the seven putative plastid pseudogenes, we searched for potential paralogs in the mitochondrial and nuclear genomes using aTRAM 2 [35]. aTRAM is a method for iterative, targeted assembly that implements commonly used *de novo* assembly modules on a reduced read set that has sequence homology with a seed sequence. Seed sequences were derived from the CDS sequence of the closest identified relative among our taxa, *Leptarrhena pyrolifolia* (D. Don) Ser. Ten iterations were used per assembly, and the assembler used was SPAdes v. 3.13.0 [36]; other options correspond to defaults. For these analyses, we extracted matching reads from the full *Saniculiphyllum* dataset (∼180,000,000 reads).

### Phylogenetics

We conducted a phylogenetic analysis both to reassess the relationships of *Saniculiphyllum* [8–10], and to assess rates of plastid substitution in a phylogenetic context. We analyzed the single-copy plastid sequence from each genome (i.e., with one copy of the inverted repeat) and ran phylogenetic analyses in RAxML v. 8.2.10 [37] under a GTR-Γ model with 1000 bootstrap replicates. Sites were partitioned as either coding (exonic protein-coding, rDNA, and tRNA) or non-coding. For this analysis, we sampled 22 further previously reported plastid genomes (Supplementary Table S1), as well as generating a plastid genome assembly from previously reported short read data from *Saxifraga granulata* L. ([38]; SRA accession SRX665162), all chosen to represent phylogenetic diversity in Saxifragales, for a total of 40 taxa. 12/16 families were sampled, including complete representation of the Saxifragaceae alliance; the plastid of the parasitic family Cynomoriaceae has been sequenced, but this was deliberately excluded as it is on an extremely long branch [39]. Saxifragaceae sampling covers 8/10 clades recognized in [9]. Tree rooting follows [10].

For the paralog search in aTRAM, we placed recovered sequences in a phylogenetic context by extracting plastid sequences for each gene from the plastid genome alignment, trimming to the extent of chloroplast gene sequences and removing ambiguously aligned regions, and removing any sequences with fewer than 200 bp remaining after these steps. We then built individual gene trees following the RAxML methods above.

### Tests for selection

For the seven loci with variation patterns suggesting putative pseudogenes, we tested for the presence of relaxed selection in *Saniculiphyllum* plastid gene copies via *ω* (dN/dS) ratios in PAML [40]. Specifically, we used a model comparison approach to ask whether the *Saniculiphyllum* branch experienced a different selection regime compared to its immediately ancestral branch; that is, whether there was a shift in selective regimes specific to this lineage. We built two models for each gene tree: a full model allowing *ω* to vary across all branches, and a constrained model where *Saniculiphyllum* was required to have the same *ω* as the branch immediately ancestral to it. We used a likelihood ratio test to determine whether the constrained model could be rejected (= a shift in selective regime along this phylogenetic branch). Since multiple tests were executed, these were corrected by the Hochberg method [41].

## Acknowledgments

D. Soltis and G. Wong are thanked for facilitating access to pilot short read data in connection with the 10KP project. J. Nelson, J. Xiang, and J.V. Freudenstein are thanked for providing DNA materials; J. Ginori assisted with testing early assembly runs, and the late M. Whitten advised extensively on DNA extraction protocols.

## Availability of data and materials

The datasets supporting the conclusions of this article are available at Dryad (alignments, partition files, and tree topologies; https://doi.org/10.5061/dryad.mgqnk98vt), and at GenBank (accession numbers in Table 2). Supplemental figures are available in Additional File 1.

## Ethics approval and consent to participate

The authors have complied with all relevant institutional, national and international guidelines in collecting biological materials for this study.

## Consent for publication

Not applicable.

## Author contributions

R.A.F. conceived the study; R.A.F. and N.S. performed analyses; B.T.S., C.-L. X., and R.P.G consulted on analyses and interpretation; R.A.F. wrote the first manuscript draft; and all authors contributed to the final manuscript draft.

