## Supplemental tables and figures for "Degradation of key photosynthetic genes in the critically endangered semi-aquatic flowering plant *Saniculiphyllum guangxiense* (Saxifragaceae)"

**Table S1.** Summary of chloroplast genome sequences downloaded from GenBank for phylogenetic analyses.

| **Species** | **Genbank accession** |
|---|---|
| *Bergenia scopulosa* | NC_036061 |
| *Cercidiphyllum japonicum* | NC_037940 |
| *Chrysosplenium aureobracteatum* | NC_039740 |
| *Chunia bucklandioides* | NC_041163 |
| *Corylopsis coreana* | NC_040141 |
| *Daphniphyllum oldhamii* | NC_037883 |
| *Fortunearia sinensis* | NC_041487 |
| *Hamamelis mollis* | NC_037881 |
| *Itea chinensis* | MH191391 |
| *Liquidambar formosana* | NC_023092 |
| *Loropetalum subcordatum* | NC_037694 |
| *Mukdenia rossii* | NC_037495 |
| *Myriophyllum spicatum* | NC_037885 |
| *Oresitrophe rupifraga* | NC_037514 |

| | |
|---|---|
| *Paeonia brownii* | NC_037880 |
| *Paeonia delavayi* | NC_035718 |
| *Parrotia subaequalis* | NC_037243 |
| *Penthorum chinense* | NC_023086 |
| *Phedimus kamtschaticus* | NC_037946 |
| *Phedimus takesimensis* | NC_026065 |
| *Rhodiola rosea* | NC_041671 |
| *Ribes fasciculatum* var. *chinense* | MH191388 |
| *Saxifraga granulata* | XXX |
| *Saxifraga stolonifera* | NC_037882 |
| *Sedum oryzifolium* | NC_027837 |
| *Sedum sarmentosum* | NC_023085 |
| *Sinowilsonia henryi* | MF497447 |

**Figure S1.** ML gene phylogeny of *ccsA*, showing the phylogenetic placement of *Saniculiphyllum* paralogs (bold) among plastid orthologs. The *Saniculiphyllum* plastid copy is marked ***. Branch labels represent bootstrap frequencies; those below 50 are not plotted.

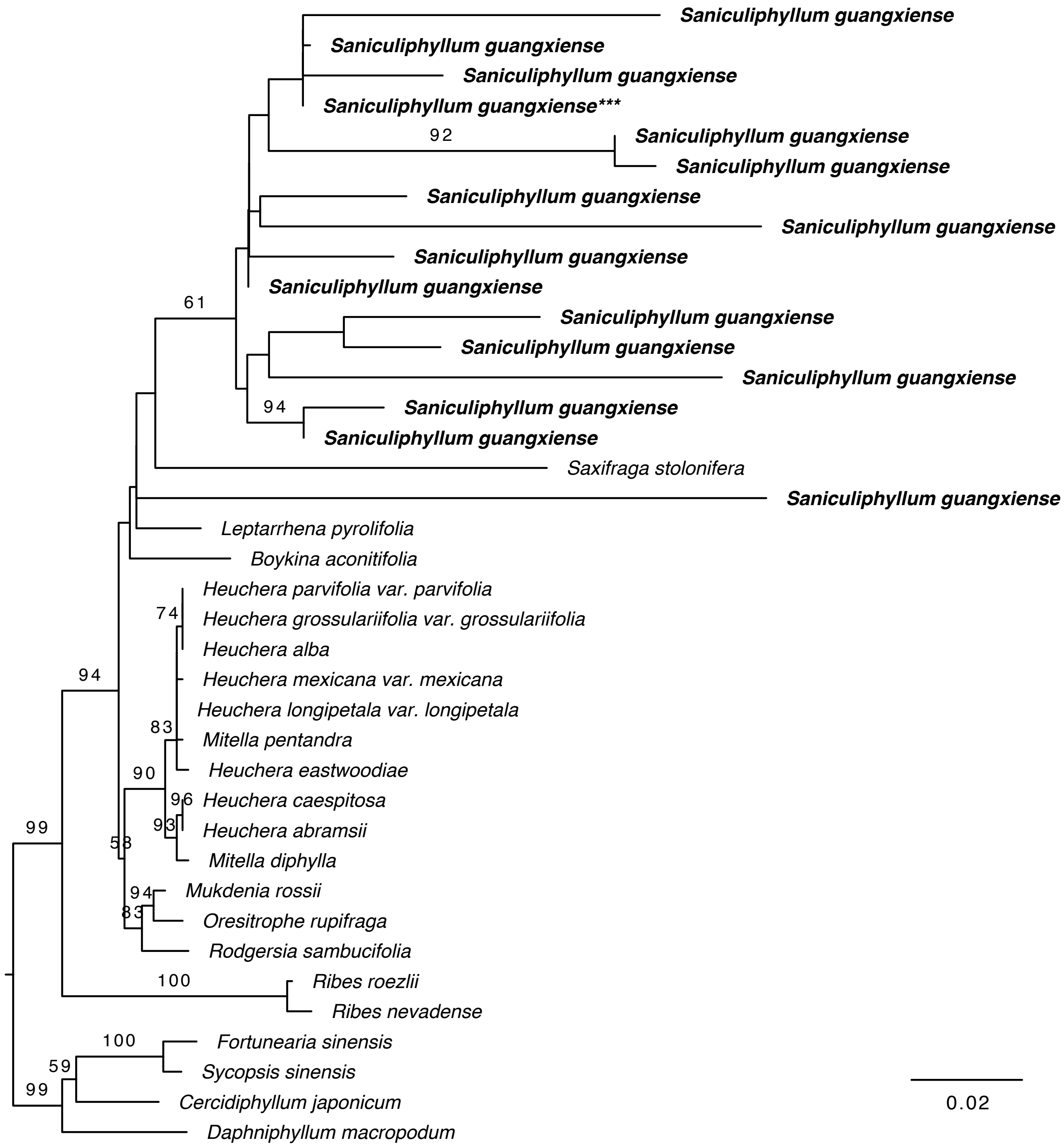

**Figure S2.** ML gene phylogeny of *cemA*, showing the phylogenetic placement of *Saniculiphyllum* paralogs (bold) among plastid orthologs. The *Saniculiphyllum* plastid copy is marked ***. Branch labels represent bootstrap frequencies; those below 50 are not plotted.

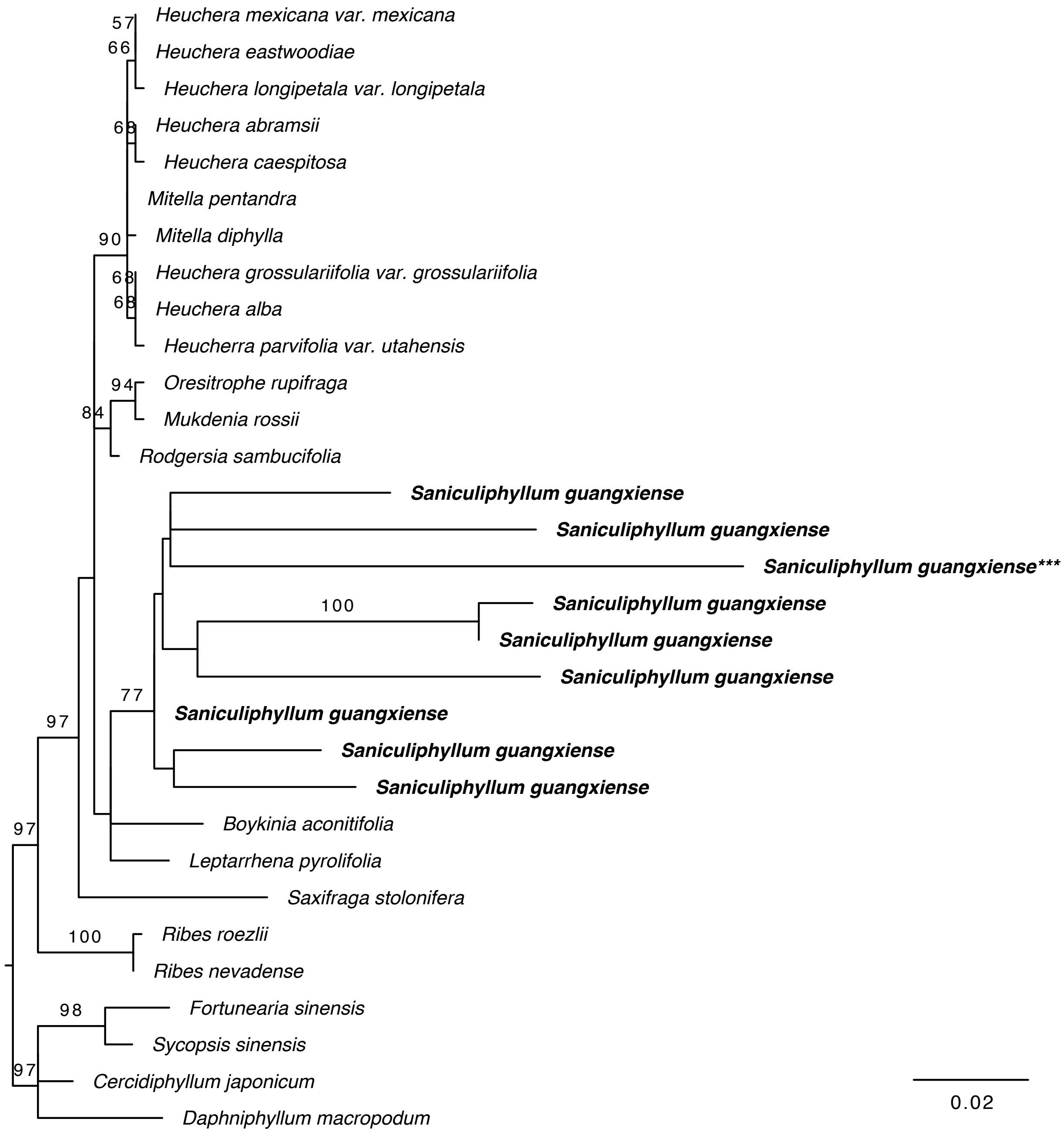

**Figure S3.** ML gene phylogeny of *ndhA*, showing the phylogenetic placement of *Saniculiphyllum* paralogs (bold) among plastid orthologs. The *Saniculiphyllum* plastid copy is marked ***. Branch labels represent bootstrap frequencies; those below 50 are not plotted.

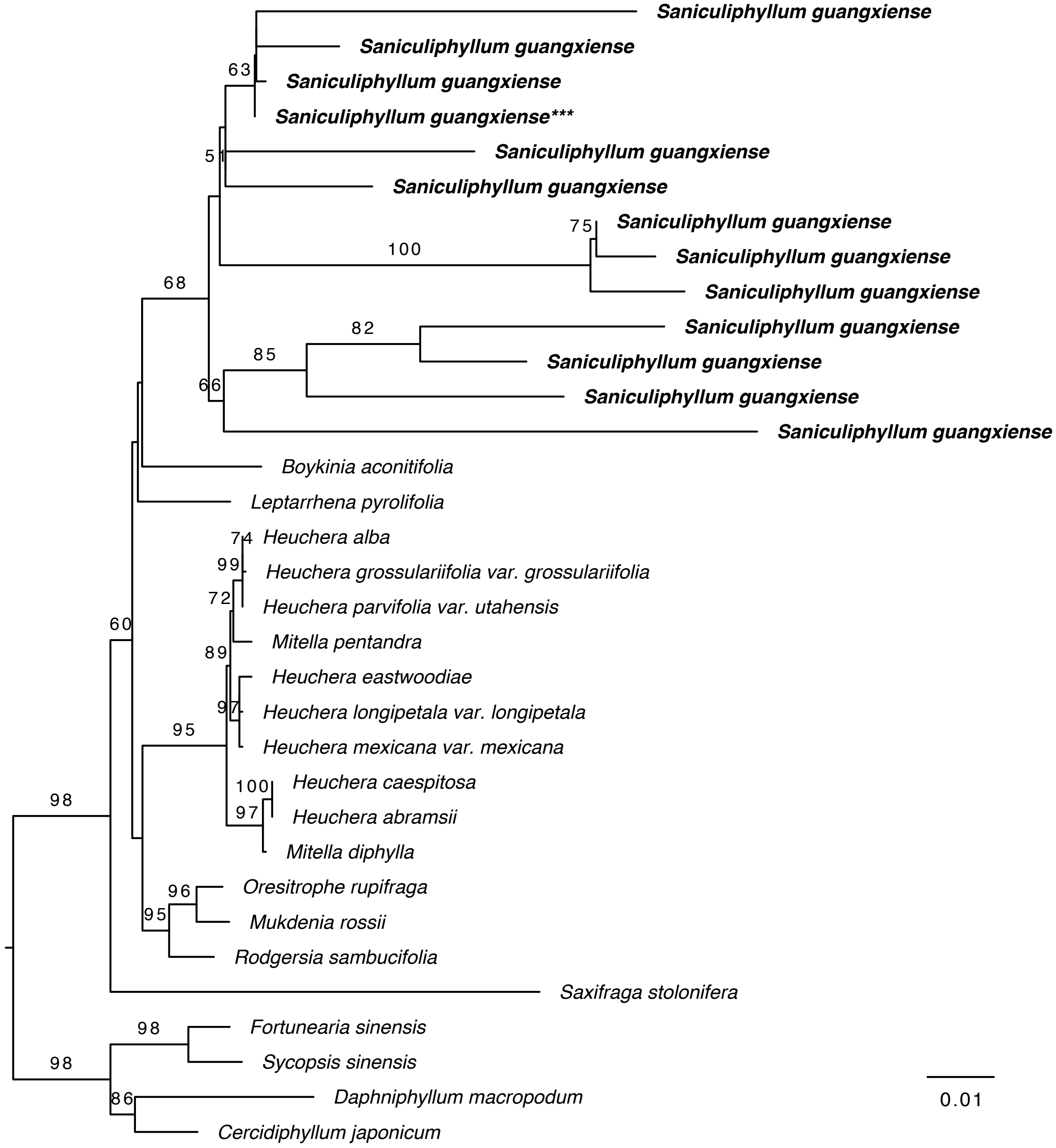

**Figure S4.** ML gene phylogeny of *ndhB*, showing the phylogenetic placement of *Saniculiphyllum* paralogs (bold) among plastid orthologs. The *Saniculiphyllum* plastid copy is marked ***. Branch labels represent bootstrap frequencies; those below 50 are not plotted.

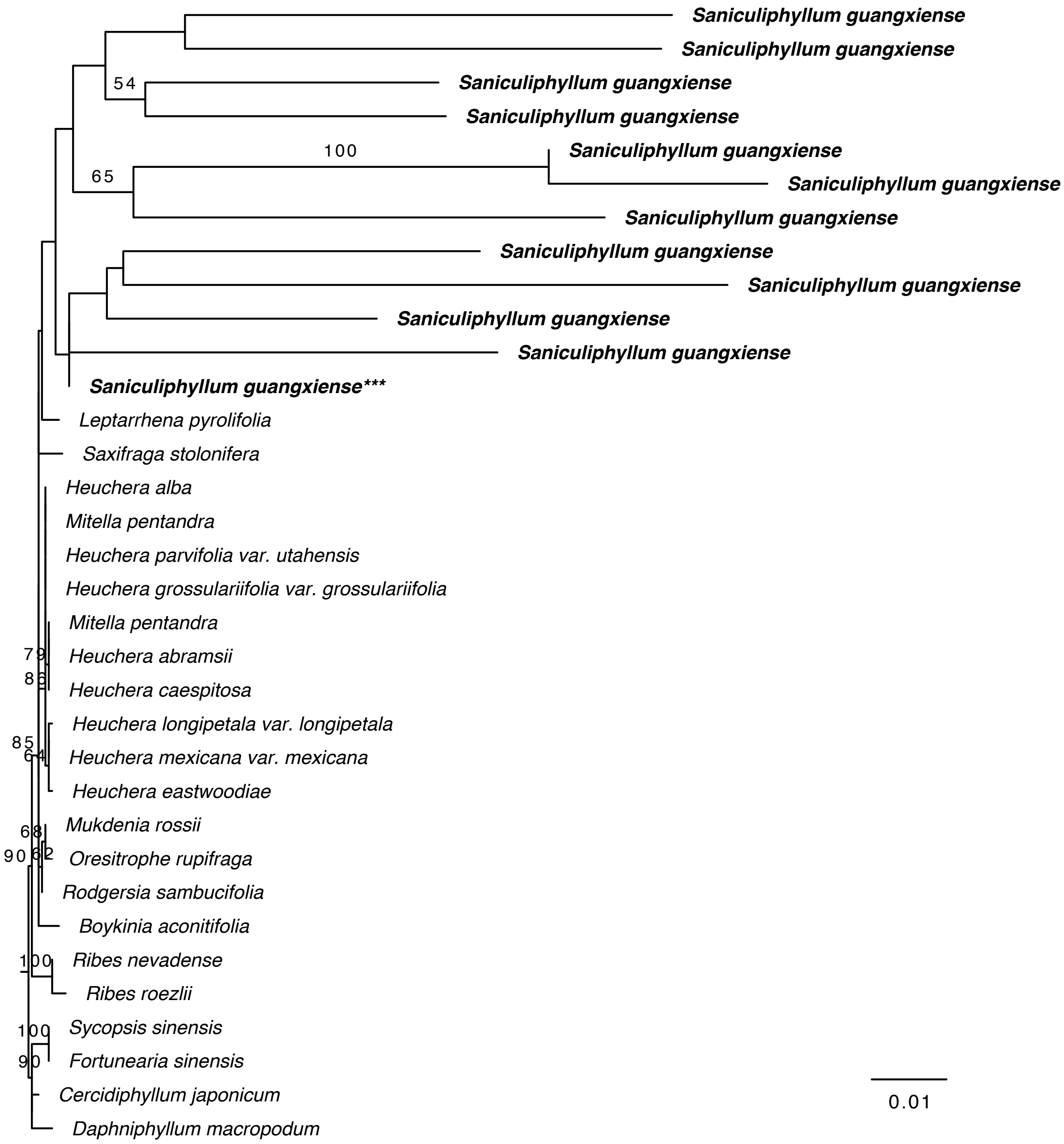

**Figure S5.** ML gene phylogeny of *ndhD*, showing the phylogenetic placement of *Saniculiphyllum* paralogs (bold) among plastid orthologs. The *Saniculiphyllum* plastid copy is marked ***. Branch labels represent bootstrap frequencies; those below 50 are not plotted.

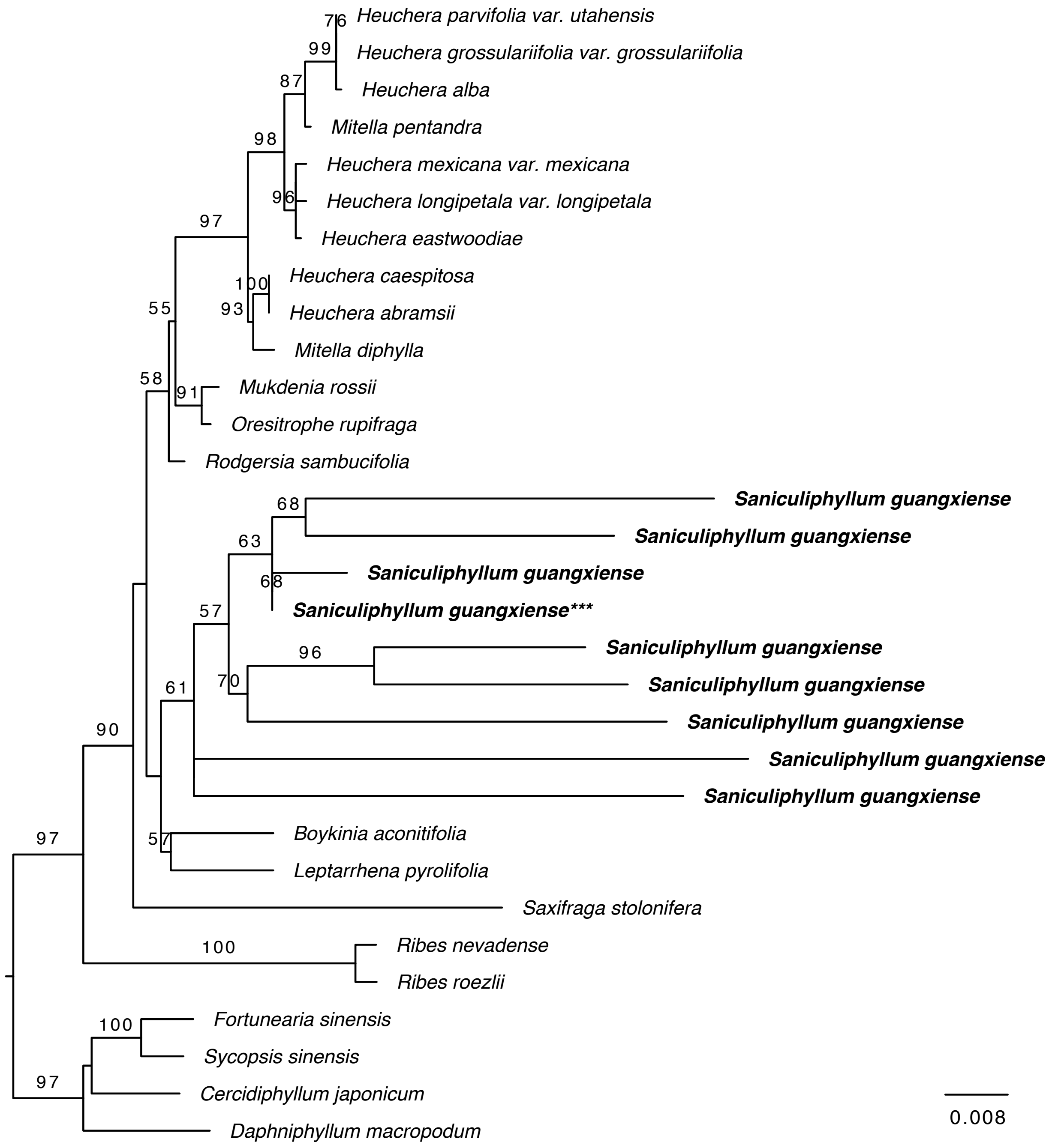

**Figure S6.** ML gene phylogeny of *ndhF*, showing the phylogenetic placement of *Saniculiphyllum* paralogs (bold) among plastid orthologs. The *Saniculiphyllum* plastid copy is marked ***. Branch labels represent bootstrap frequencies; those below 50 are not plotted.

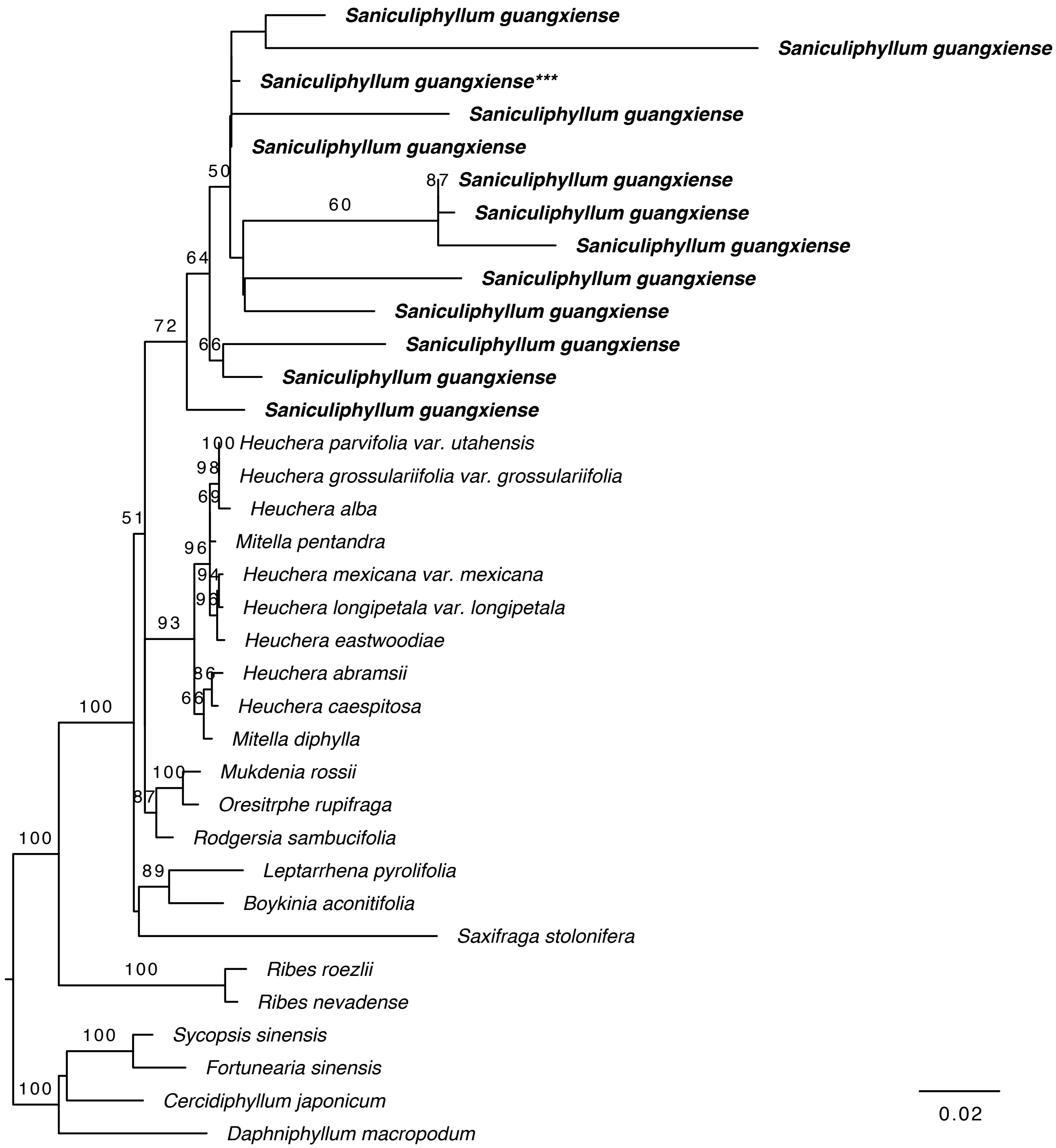

**Figure S7.** ML gene phylogeny of *ndhK*, showing the phylogenetic placement of *Saniculiphyllum* paralogs (bold) among plastid orthologs. The *Saniculiphyllum* plastid copy is marked ***. Branch labels represent bootstrap frequencies; those below 50 are not plotted.

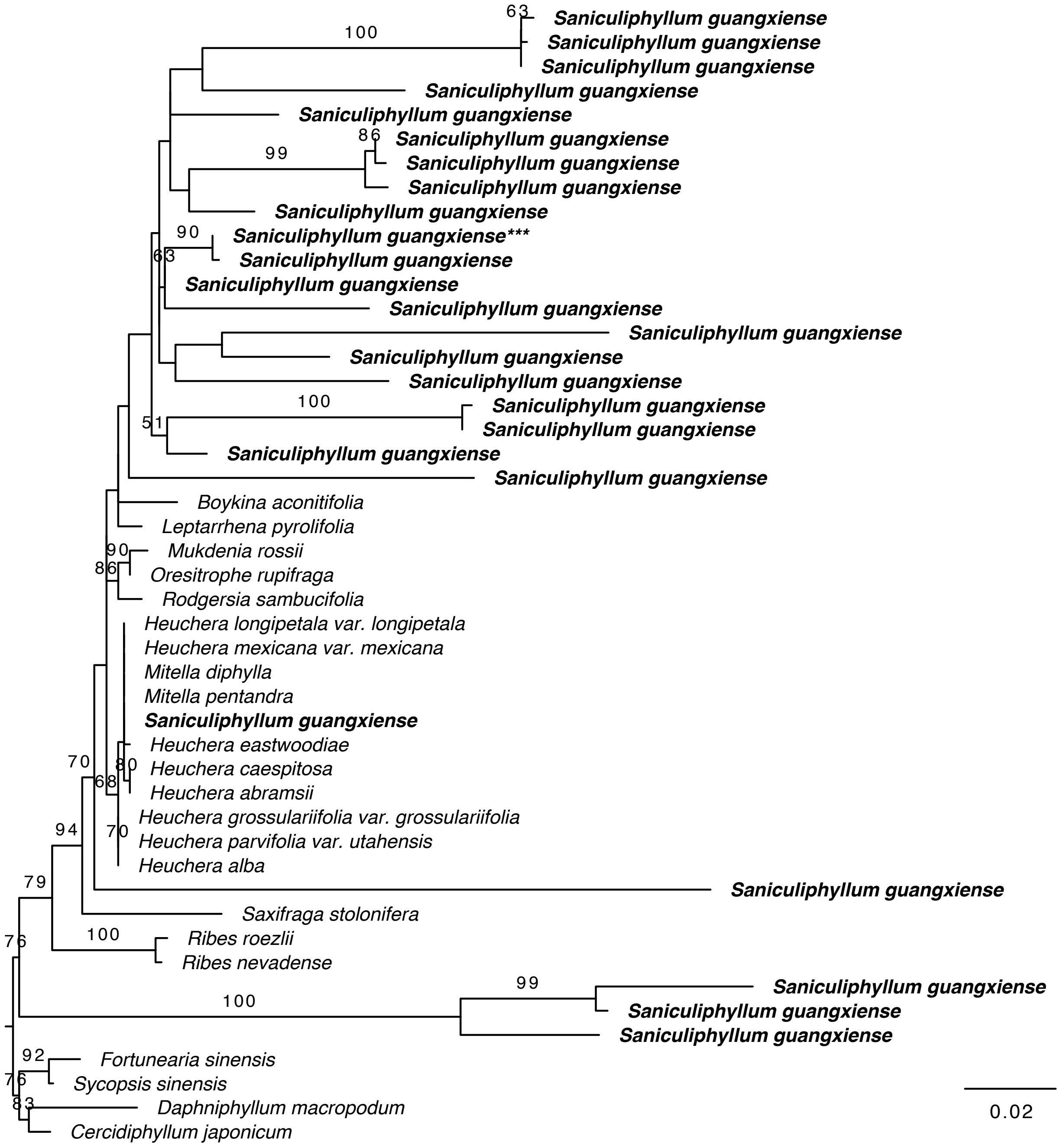
